# A series of five population-specific Indian brain templates and atlases spanning ages 6 to 60 years

**DOI:** 10.1101/2020.05.08.077172

**Authors:** Bharath Holla, Paul A. Taylor, Daniel R. Glen, John A. Lee, Nilakshi Vaidya, Urvakhsh Meherwan Mehta, Ganesan Venkatasubramanian, Pramod Pal, Jitender Saini, Naren P. Rao, Chirag Ahuja, Rebecca Kuriyan, Murali Krishna, Debashish Basu, Kartik Kalyanram, Amit Chakrabarti, Dimitri Papadopoulos Orfanos, Gareth J. Barker, Robert W. Cox, Gunter Schumann, Rose Dawn Bharath, Vivek Benegal

## Abstract

Anatomical brain templates are commonly used as references in neurological MRI studies, for bringing data into a common space for group-level statistics and coordinate reporting. Given the inherent variability in brain morphology across age and geography, it is important to have templates that are as representative as possible for both age and population. A representative-template increases the accuracy of alignment, decreases distortions as well as potential biases in final coordinate reports. In this study, we developed and validated a new set of T1w Indian brain templates (IBT) from a large number of brain scans (total n=466) acquired across different locations and multiple 3T MRI scanners in India. A new tool in AFNI, make_template_dask.py, was created to efficiently make five age-specific IBTs (ages 6-60 years) as well as maximum probability map (MPM) atlases for each template; for each age-group’s template-atlas pair, there is both a “population-average” and a “typical” version. Validation experiments on an independent Indian structural and functional-MRI dataset show the appropriateness of IBTs for spatial normalization of Indian brains. The results indicate significant structural differences when comparing the IBTs and MNI template, with these differences being maximal along the Anterior-Posterior and Inferior-Superior axes, but minimal Left-Right. For each age-group, the MPM brain atlases provide reasonably good representation of the native-space volumes in the IBT space, except in a few regions with high inter-subject variability. These findings provide evidence to support the use of age and population-specific templates in human brain mapping studies. This dataset is made publicly available (https://hollabharath.github.io/IndiaBrainTemplates).

**Highlights:**

1. A new set of age-specific T1w Indian brain templates for ages 6-60 yr are developed and validated.
2. A new AFNI tool, make_template_dask.py, for the creation of group-based templates.
3. Maximum probability map atlases are also provided for each template.
4. Results indicate the appropriateness of Indian templates for spatial normalization of Indian brains

## 1 Introduction

The shape, size and volume of the human brain is highly variable across individuals, as well as across age, gender and geographical location or ethnicity. This fact is of prime importance in neuroimaging group studies, where the brains of all subjects are typically aligned to a single template space for data analysis and for the reporting of findings where analogous anatomical structures are mapped on to the same coordinate location across the subjects. A brain template provides a standard 3D coordinate frame to combine and/or compare data from many subjects, across different imaging modalities, structural or functional and even different laboratories around the world. The properties of the template (size, shape, tissue contrast, etc.) directly affect the quality of alignment.

An early brain atlas was constructed by Talairach and Tournoux [1988] from a post mortem brain of one 60-year-old French woman, introducing the concepts of coordinate system and spatial transformation to brain imaging. However, using a single subject brain as a template introduces several idiosyncrasies, as it does not account for groupwide anatomical variability, asymmetry, age-related differences, etc. In order to address some of these issues, a subsequent initiative from the Montreal Neurological Institute (MNI) resulted in a statistical brain template (MNI-305) using 305 young right-handed subjects [Evans et al., 1993]. While this composite template better accounted for anatomical variability, it also had relatively low tissue contrast and structural definition, which can affect the ability of alignment algorithms to provide high quality anatomical matching across a group study. In 2001, the international consortium for human brain mapping (ICBM) introduced the revised MNI-152 template [Mazziotta et al., 2001b] with better contrast and structure definition, where 152 individual brains were linearly registered to MNI305 to make an average template. The ICBM-452 template [Mazziotta et al., 2001a] included all three sites of ICBM and provided even better signal-to-noise ratio due to the nearly threefold increase in the number of subjects. These MNI templates were widely adopted by several image processing pipelines, with the associated set of coordinates known as “MNI space”. Furthermore, an unbiased non-linear average of the adult MNI152 and a pediatric template with 20-40 iterative non-linear averages has also been made available [Fonov et al., 2011]. These templates provide the advantages of retaining group representativeness of the MNI305 or MNI152 while still providing the details that are closer to those apparent in a single subject; however, their “representativeness” is limited to a fairly isolated geographic location and (typically, Western) population, even though neuroimaging studies draw from populations across the globe.

More recently, several research groups around the world have developed and validated brain templates that are representative of their (broadly) local population. Lee et al. [2005] created a set of Korean Brain templates with 78 subjects in an age range between 18 to 77 years (young template <55 years and elderly template >55 years). Additionally, Tang et al. [2010] generated a Chinese brain template of 56 subjects (mean age 24.4 years). In each case the groups demonstrated significantly reduced warp deformations and increased registration accuracy when applying these templates to studies of local populations. It should be noted that even though the templates draw from subjects within a population, there is still a large amount of inherent variability evident in the brain morphology, due to combinations of factors such as inherent structural variability, multi-ethnic composition and differences in genetic influences and environmental exposures.

The benefit of utilizing a population-representative template in the Indian context has also been recognized, with the additional need for age-specific templates due to the increasingly wide range of ages enrolled in studies. Recent attempts at developing brain templates for Indian population have tended to focus on the young adult age group (21-30 years) with relatively small [Rao et al., 2017] to modest sample sizes [Sivaswamy et al., 2019, Bhalerao et al., 2018, Pai et al., 2020], and have utilized data from a single site/scanner. Additionally, to date, whole-brain annotated reference atlases based on segmentation have not accompanied the generated templates. In this study, we present and validate a new set of brain templates that have been created from a large number of subjects from multi-site acquisitions across India, with five age ranges provided (between 6-60 years), as well as brain atlases for each template. For each age group’s template-atlas pair, there is both a “population average” and “typical” version (the latter being the individual brain which most closely matches the population average, which potentially provides higher detail as an alignment target and atlas). We present several validation tests for the accuracy and representativeness of the templates, and we also use data from separately acquired subjects to demonstrate the benefits of these templates over the existing standard MNI templates for studies on Indian cohorts.

## 2 Methods

### 2.1 Participants

The datasets used in the present study were selected retrospectively from healthy control subjects of several imaging studies, across multiple centers and different populations across India. They included imaging data from the ongoing Indian multi-site developmental cohort study, the Consortium on Vulnerability to Externalising Disorders and Addictions (cVEDA) [Sharma et al., 2020, Zhang et al., 2020] and from stored datasets contributed by researchers at the National Institute of Mental Health and Neurosciences (NIMHANS, Bengaluru, India). All of these studies were approved by the ethics review boards at the corresponding participating sites and informed consent was obtained from each participant (or from their parent, in the case of subjects below 16 years, along with participant’s written assent) with a specific request to collect, store and share anonymized data for research. Inclusion criteria included not having a personal history of prior brain injury, neurological disorder or psychiatric diagnosis. The sample was comprised of 466 subjects from a large number of states across India and acquired at multiple sites. Based on age and demographic distributions, subject datasets were divided into 5 groups: C1, late childhood (6-11 years); C2, adolescence (12-18 years); C3, young adulthood (19-25 years); C4, adulthood (26-40 years); C5, late adulthood (41-60 years). The sample size and demographic information of each cohort is summarized in Table 1.

**Table 1.** Demographic Profiles.

| Age Category | Age Description | Age in years, Mean (Range) | Sample Size $N$ (% Female) | No. States | No. Scanners |
| --- | --- | --- | --- | --- | --- |
| C1 | Late childhood | 9.3 (6 to 11) | 28 (46.43%) | 5 | 4 |
| C2 | Adolescence | 15.1 (12 to 18) | 106 (47.17%) | 9 | 5 |
| C3 | Young adulthood | 21.3 (19 to 25) | 181 (40.89%) | 15 | 5 |
| C4 | Adulthood | 31.1 (26 to 40) | 89 (42.7%) | 11 | 2 |
| C5 | Late adulthood | 52.7 (41 to 60) | 62 (43.55%) | 6 | 2 |

### 2.2 Image acquisition

T1-weighted (T1w) three-dimensional high resolution structural brain MRI scans were acquired from five 3T MRI scanners located at three different locations across India: Bengaluru (site A, C and D), Mysuru (site B) and Chandigarh (site E). The subjects belonged to several neighboring states to these locations, with wide geographical representation throughout India. As with most multisite studies, the acquisition parameters varied slightly across sites and scanners, but were generally similar, with good grey/white matter contrast with a voxel size close to 1mm isotropic; details are listed in Table 2.

**Table 2.** Acquisition parameters.

| Acq Seq | Site label | Scanner model | dx (mm) | dy (mm) | dz (mm) | TR<sup>†</sup> (ms) | TE (ms) | TI (ms) | FA (deg) | Matrix size | No. Sag | No. Subj<sup>‡</sup> |
| --- | --- | --- | --- | --- | --- | --- | --- | --- | --- | --- | --- | --- |
| 1 | A | Achieva<sup>a</sup> | 1 | 1 | 1 | 8.2 | 3.8 | 745 | 8 | 256 × 256 | 165 | 50 |
| 2 | A | Achieva<sup>a</sup> | 0.9 | 0.9 | 1 | 8.2 | 3.8 | 800 | 8 | 257 × 256 | 160 | 38 |
| 3 | B | Ingenia<sup>a</sup> | 1.2 | 1 | 1 | 6.9 | 3.2 | 725 | 9 | 256 × 256 | 170 | 29 |
| 4 | C | Ingenia<sup>a</sup> | 1 | 1 | 1 | 6.9 | 3.3 | 925 | 9 | 256 × 256 | 211 | 10 |
| 5 | D | Skyra<sup>b</sup> | 1.2 | 1 | 1 | 2300 | 3.0 | 900 | 9 | 256 × 240 | 176 | 82 |
| 6 | D | Skyra<sup>b</sup> | 1 | 1 | 1 | 1900 | 2.4 | 900 | 9 | 256 × 256 | 192 | 56 |
| 7 | D | Skyra<sup>b</sup> | 0.9 | 0.9 | 0.9 | 1600 | 2.1 | 900 | 9 | 256 × 256 | 176 | 124 |
| 8 | E | Verio<sup>b</sup> | 1.2 | 0.5 | 0.5 | 2300 | 3.0 | 900 | 9 | 512 × 480 | 176 | 77 |

Acq Seq = acquisition sequence; dx, dy, dz are voxel dimensions; TR = repetition time; TE = echo time; TI = inversion time; FA = flip angle; No. Sag = number of sagittal slices.
<sup>a</sup>Philips, 3T. <sup>b</sup>Siemens, 3T. <sup>‡</sup>This is the final number of subjects included in final templates (total = 466), after all steps of QC and subject removal. <sup>†</sup>The TR for 3D scans such as these is defined differently between Philips and Siemens scanners, with the relationship being $TR_{\text{Philips}} \approx (TR_{\text{Siemens}} - TI)/(No. \text{ Sag})$ .

### 2.3 Data Preprocessing and Initial Quality Assurance

This processing primarily used programs in the AFNI (v19.0.20) [Cox, 1996] and FreeSurfer (v6.0) [Fischl, 2012] neuroimaging toolboxes, as well as the “dask” scheduling tool in Python developed by the Dask Development Team [2016]. Unless otherwise noted, programs named here are contained within the AFNI distribution. The following processing steps are shown schematically in Figure 1, in the first column.

**Figure 1.**
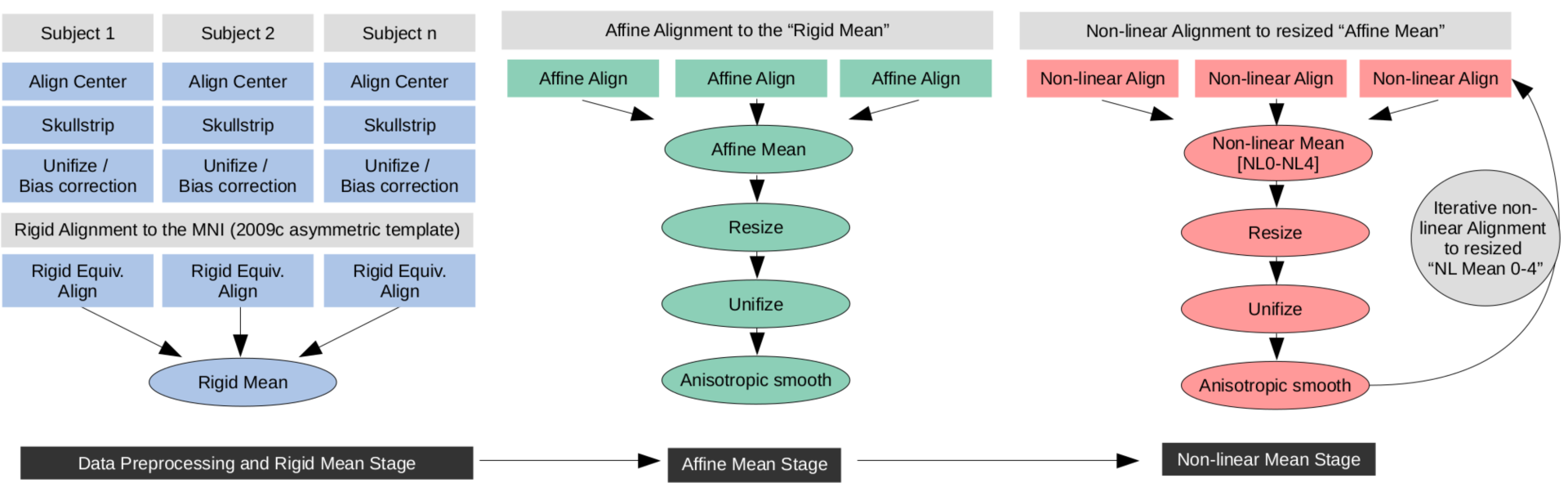
Schematic representation of the steps involved in the Dask pipeline (make_template_-dask.py) for generating population-average brain templates.

Datasets were first processed using AFNI’s “fat_proc_convert_dcm_anat”. Using this, DICOMs were converted to NIFTI files using dcm2niix_afni (the AFNI-distributed version of dcm2niix [Li et al., 2016]). For uniformity and initialization, with this tool, they were also given the same orientation (RAI), and the physical coordinate origin was placed at the volume’s center of mass (to simplify later alignments).

Next, “fat_proc_axialize_anat” was applied to reduce the variance in the spatial orientation of brains for later alignment and for practical considerations of further processing steps, as described here. Each volume was affinely registered to a reference anatomical template (MNI ICBM 152 T1w) that had previously been AC-PC aligned; alignment included an additional weight mask to emphasize subcortical structure alignment (e.g., AC-PC structures), and only the solid-body parameters of the alignment were applied, so that no changes in shape were incurred. Because datasets had been acquired with varied spatial resolution and FOV (see Table 2), the datasets were resampled (using a high-order sinc function, to minimize smoothing) to the grid of the reference base of 1mm isotropic voxels.

All datasets were visually and systematically checked for quality of both data and registration using the QC image montages that were automatically generated by the previous program. T1w volumes with noticeable ringing or other artifact (e.g., due to subject motion or dicom reconstruction errors) were noted and removed from further analyses. T1w volumes with any incidental findings (for example, large ventricles, cavum septum pellucidum) were also removed.

FreeSurfer’s “recon-all” [Fischl, 2012] was run on each T1w data set to estimate surfaces, parcellation and segmentation maps. AFNI’s “@SUMA_Make_Spec_FS” was then run to convert the FreeSurfer output to NIFTI files and to generate standard meshes of the surface in formats usable by AFNI and SUMA. Additionally, @SUMA_Make_Spec_FS subdivides the FreeSurfer parcellations into tissue types such as gray matter (GM), white matter (WM), cerebrospinal fluid (CSF), ventricle, etc. This was followed by visual inspection of parcellation maps overlaid on anatomical volumes.

Next, a whole brain mask of each anatomical volume was created. In several cases, the skullstripped brain volumes output by recon-all (brain_mask.nii) included large amounts of non-brain material (skull, dura, face, etc.), and so an alternative mask was generated using only the ROIs comprising the parcellation and segmentation maps. For each subject, a whole brain mask was generated by: first making a preliminary mask from all of the ROIs identified by recon-all; then inflating that premask by 3 voxels; and finally shrinking the result by two voxels (thus filling in any holes inside the brain mask and smoothing the outer edges). This produced whole brain masks that were uniformly specific to each subject’s intracranial volume.

Finally, AFNI’s 3dUnifize was run on each T1w volume in order to reduce the intensity inhomogeneity (e.g., due to the bias field) and to normalize the intensity of tissues within the volume. This ensures that each subject’s brain, which had been acquired on different scanners with potentially different scalings, would have equal weight when averaging (e.g., WM is scaled to approximately a value of 1000 in each brain, and similarly for other tissues), and also reduces the risk of a bright outlier region driving poor alignment.

### 2.4 Mean template generation

After the above pre-processing steps and QC, the following templatizing algorithm was applied for each cohort (C1-5) separately. The general procedure was to alternate between alignment to a reference base (with increasingly higher order of refinement) and averaging the aligned brains to generate a new reference base for the subsequent iteration. In this way one can generate a cohort mean template of successively greater specificity and detail; after several iterations, the alignment essentially converges (i.e., additional refinement becomes negligible) and is halted. Warps were generated and saved at each step. The final nonlinear warps and affine transformations were concatenated for each subject at the end in order to generate the final group average template. These steps are also included in the schematic Figure 1, in the first column (bottom) and second and third columns.

The first level of alignment was made from each anatomical in the cohort to the MNI ICBM-152 T1w template using a 6 degree of freedom (DF) rigid body equivalent registration, meaning a full affine transformation was computed, but only the rigid components were extracted and applied. The average of all subjects’ brains, rigidly aligned to the initial template, was used to create a single average volume “mean-rigid”; here and at each alignment stage, a cohort standard deviation map was also created, to highlight locations of relatively high and low variability. That stage’s average volume was then used as a base for the next stage of alignment for each subject, using a 12 DF linear affine registration, and with the results averaged to create the next base “mean-affine”. For these alignments, AFNI’s “lpa” cost function (absolute value of local Pearson correlation) [Saad et al., 2009] was used for high quality alignment of features between volumes of similar contrast. The cost function computes the absolute value of the Pearson correlation between the volume and the current template in patches of the volume at a time.

As a practical consideration, we note that lower level alignments such as these have a general property of producing a smoothed brain, which has the additional effect of increasing the apparent size of the base dataset (i.e., the edge is blurred outward). Therefore, in these initial levels we added a step to control the overall volume of the template. We calculated the mean intracranial volume (ICV) of all the subjects in the cohort *V*_coh_, and then calculated the volume of the initial mean-affine brain mask *V*_aff_. The volume ratio *r*_vol_ = *V*_coh_*/V*_aff_ was calculated, and each of the three dimensions of the mean-affine volume were scaled down by the appropriate length scaling factor 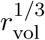. In this way, the final volume of the templating process retained a representative size for the cohort.

The next alignment stages were comprised of nonlinear registration using AFNI’s 3dQwarp [Cox and Glen, 2013]. At each successive level the nonlinear alignment was performed to an increasingly higher refinement, resulting in mean volumes of greater detail. Specifically, nonlinear alignment at each stage was implemented to create mean templates as follows (A-E), using 3dQwarp’s default “pcl” (Pearson correlation, clipped) cost function to reduce the effects of any outlier values (and unless otherwise specified, applying a 3D Gaussian blur):

A. **mean-NL0**: after registering to mean-affine with a minimum patch size of 101 mm and blurring of 0 mm (base) and 9 mm (source);
B. **mean-NL1**: after registering to mean-NL0 with a minimum patch size of 49 mm and blurring of 1 mm (base) and 6 mm (source);
C. **mean-NL2**: after registering to mean-NL1 with a minimum patch size of 23 mm and blurring of 0 mm (base) and 4 mm (source);
D. **mean-NL3**: after registering to mean-NL2 with a minimum patch size of 13 mm and blurring of 0 mm (base) and 2 mm median filter (source);
E. **mean-NL4**: after registering to mean-NL3 with a minimum patch size of 9 mm and blurring of 0 mm (base) and 2 mm median filter (source).

Each mean-NL* volume was resized in the same manner as the initial stages, although the correction factors were much smaller here. Additionally, each mean-NL* volume was anisotropically smoothed (preserving edges within the volume, for detail) using 3danisosmooth, in order to sharpen its contrast for subsequent alignments.

The mean-NL4 volume became the final group mean template for each cohort, as in all cases results appeared to have essentially converged after this number of step. The coordinate system of this mean volume defines the template space for that age group, and is labelled “IBT_C1”, “IBT_C2”, etc.

### 2.5 “Typical” subject template generation

We used the following approach to find the maximally representative individual brain for the mean template from the underlying cohort, in order to generate an additional “typical” template for that space, in complement to the mean template.

To find the most typical subject for the mean template quantitatively, the lpa cost function value from aligning each subject’s anatomical to the final mean-NL4 was compared across the group; that is, the degree of similarity of each subject’s aligned volume to the mean template base was compared across the cohort. The individual brain in that mean template space with the lowest cost function value was selected to be the “typical template” brain. Alignment results were also visually verified for each typical template. We note that the typical template volume uses the same coordinate system as the mean template, and thus no additional “coordinate space” is created in this process.

### 2.6 Atlas generation for mean and typical templates

For each cohort, atlases were generated for each of the mean and typical templates based on FreeSurfer parcellation and segmentation maps^1^. By default, recon-all produces two maps of ROIs (including both cortical and subcortical GM, WM, ventricles, etc.): the “2000” map, using the Desikan-Killiany Atlas [Desikan et al., 2006] and the “2009” map, using the Destrieux Atlas [Destrieux et al., 2010]. Each of these maps was used to create a “2000” and “2009” atlas for each template.

For the mean template, maximum probability map (MPM) atlases were reconstructed as follows. The FreeSurfer parcellations for each subject were transformed to the IBT space using the warps created during the template creation process (and “nearest neighbor” interpolation, to preserve ROI identity). For a given parcellation, the fraction of overlap of a given ROI at each voxel in the template was computed. That overlap fraction is essentially the probability of a region to be mapped to that voxel. In this way, an MPM atlas was created for each of the 2000 and 2009 parcellations, labelled “IBT_C1_MPM_2000”, “IBT_C1_MPM_2009”, etc. The value of each voxel’s maximum probability was also kept and stored in a map, for reference and validation. Locations with max probability near 1 show greatest uniformity across group, and locations with lower values show greater variability.

For each typical template volume, atlases based on the 2000 and 2009 FreeSurfer parcellation were also created. First, the parcellations from original subject space were mapped to the individual template space. Then, each parcellation was passed through a modal smoothing process using 3dLocalstat: for each voxel in the atlas, its value was reassigned to the mode of its NN=1 neighborhood (i.e., among “facewise” neighbors, so within a 7 voxel neighborhood). In this way the final atlas parcellation was slightly regularized, in order to reduce the effects of resampling to the template space. A typical brain atlas was created from each of the 2000 and 2009 parcellations, labelled “IBT_C1_TYP_2000”, “IBT_C1_TYP_2009”, etc.

### 2.7 Validation and tests

The fractional volumes of each ROI in the MPM atlases were checked for being representative of each cohort. For this we calculated the logarithm of the relative volume ratio of each ROI:

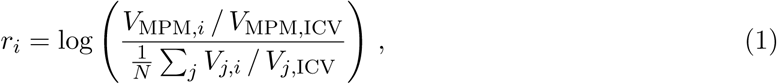

where the numerator is the fractional volume of a given *i*th ROI in the MPM (i.e., volume of the ROI divided by that template’s ICV), and the denominator is the fractional volume of that *i*th ROI averaged across all *N* subjects (i.e., for each *j*th subject, volume of the ROI divided by the subject’s ICV, in native space). Thus, *r*_*i*_ values close to 0 reflect high similarity of the MPM ROI to the cohort mean, and negative or positive values reflect a relative compression or expansion, respectively, of the MPM ROI relative to that for a particular cohort.

In order to quantify the inter-subject brain morphological variability for participants in each ageband, we calculated a region-wise mean deformation value (mDV) from the deformation warp fields generated during non-linear registration to the age-specific IBT. For this, the absolute warp value was summed across all three axes (L1-norm) and averaged across all the voxels within each ROI in the age-specific MPM atlas. A larger mDV indicates greater inter-subject brain morphological variability.

To examine the utility of the IBTs on a real, representative dataset, a separate sample of Indian population data was included for validation and testing purposes. For each cohort, the validation group (“V1”, matched with cohort C1; “V2”, matched with cohort C2; etc.) comprised 20 subjects within the corresponding age range. The T1w and resting state functional MRI (rs-fMRI) data acquisition information and demographics of these additional groups are provided in supplementary text. For each IBT, in comparison to the MNI ICBM-152 template, the following validation tests were conducted using the T1w and resting functional data.

We first used the deformation field to characterize the difference between the two templates (IBT vs MNI). For each subject in the validation cohort, we calculated the absolute amount of displacement needed to move a voxel location from native space to the target in the new age-specific IBT and the standard MNI ICBM-152 templates, for non-linear registration. A median absolute distance along each axis (LR = left-right; PA = posterior-anterior; IS = inferior-superior) was calculated from the dimensional deformation field in each voxel. The median absolute distances when warping to MNI and cohort-specific IBT along each axis were compared using a paired sample Wilcoxon’s signed-ranks test.

Finally, the practical benefits of using the IBT as reference volume for FMRI alignment were investigated by processing resting state FMRI data from age-specific validation cohorts using the same pipeline twice: once with the IBT, and once with the standard MNI template. AFNI’s afni_proc.py command was used to generate the full fMRI processing pipeline and the exact command is provided in the supplementary text. We used AFNI’s 3dReHo [Taylor and Saad, 2013] to calculate a common resting state FMRI parameter, ReHo (region homogeneity, which is Kendall’s Coefficient of Concordance, W, in statistics [Kendall and Smith, 1939, Zang et al., 2004]), within each atlas ROI for the data in each of the IBT and MNI spaces (as per template-specific Desikan-Killiany Atlas, which exists in both spaces). We then performed a paired t-test comparison on the ROI-ReHo values, in order to compare ReHo values between template space targets. In the current pair-wise comparisons, a greater ReHo would indicate greater temporal coherence of BOLD time series, likely due improvement in overall alignment across subjects within each ROI.

## 3 Results

The first part of the output consists of both “population average” and “typical” Indian brain templates for five specific age-ranges: late-childhood (C1), adolescence (C2), young adulthood (C3), adulthood (C4) and late adulthood (C5) [see Table 1 for the age-ranges]. The second part of the output is a set four IBT atlases (IBTAs) for each age range: both an MPM and a typical subject version of each of the Desikan-Killiany (FreeSurfer’s “2000”) and Destrieux (FreeSurfer’s “2009”) atlases.

Figure 2 shows an example of the successive stages in the creation of the C1 IBT. Throughout the refinement, details become progressively clearer, with tissue contrast and feature identification increasing. Additionally, the variance decreases in the gray and white tissues with each stage. The contrast-to-noise ratio (CNR) between GM and WM improved through the successive stages in all the template age-groups (see Supplementary Figure S1).

**Figure 2.**
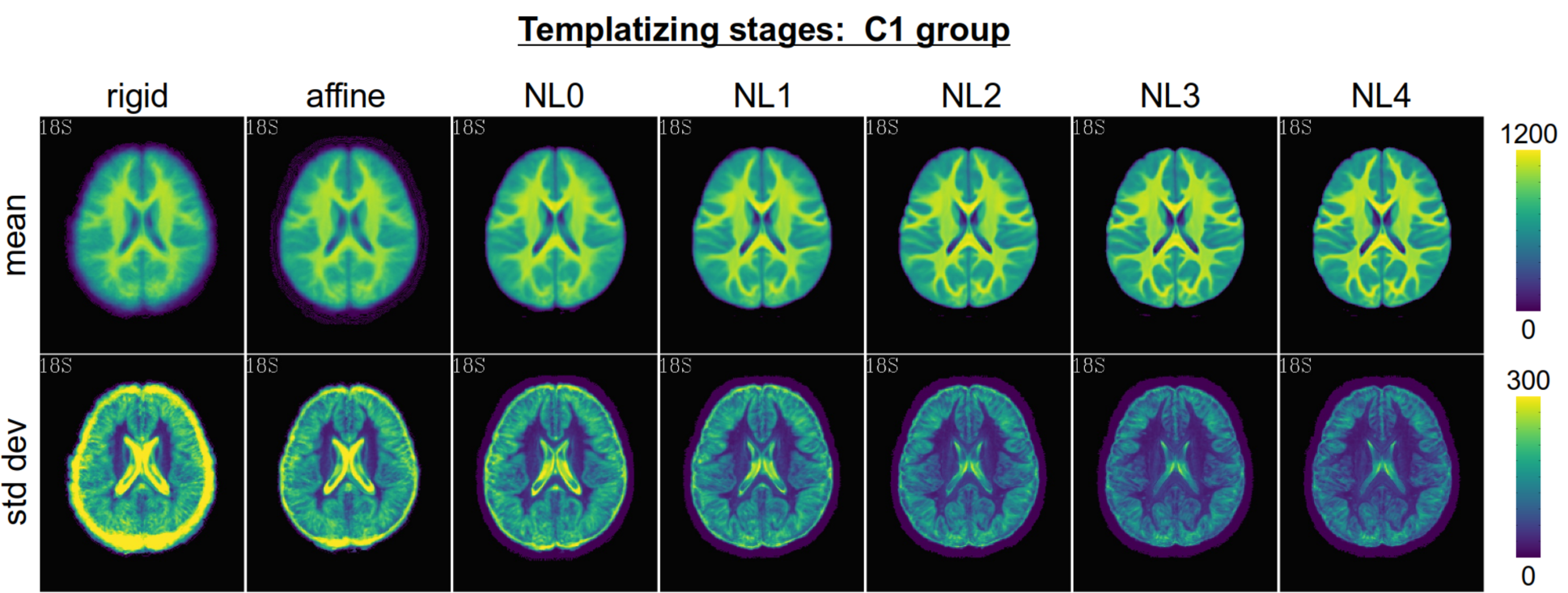
Axial slices of mean (top row) and standard deviation (bottom row) maps through successive stages of the templatizing algorithm (first stage at the left) for the C1 age-band. Note that the mean and standard deviation maps have separate scales, to show details more clearly in each.

Figure 3 shows an example of the IBT and IBTA outputs for the C3 group, displaying multiple slices in sagittal, coronal and axial views; in all cases, the population average template is underlayed. The top row shows a size comparison with the overlaid MNI template (shows as edges). In the second row, the “typical” template version is overlaid translucently, showing the very high degree of structural similarity between the two template versions. The bottom two rows show the MPM 2000 and 2009 IBTAs. Similar outputs for other age groups are provided in the Supplementary Information, in Figures S2-S6.

**Figure 3.**
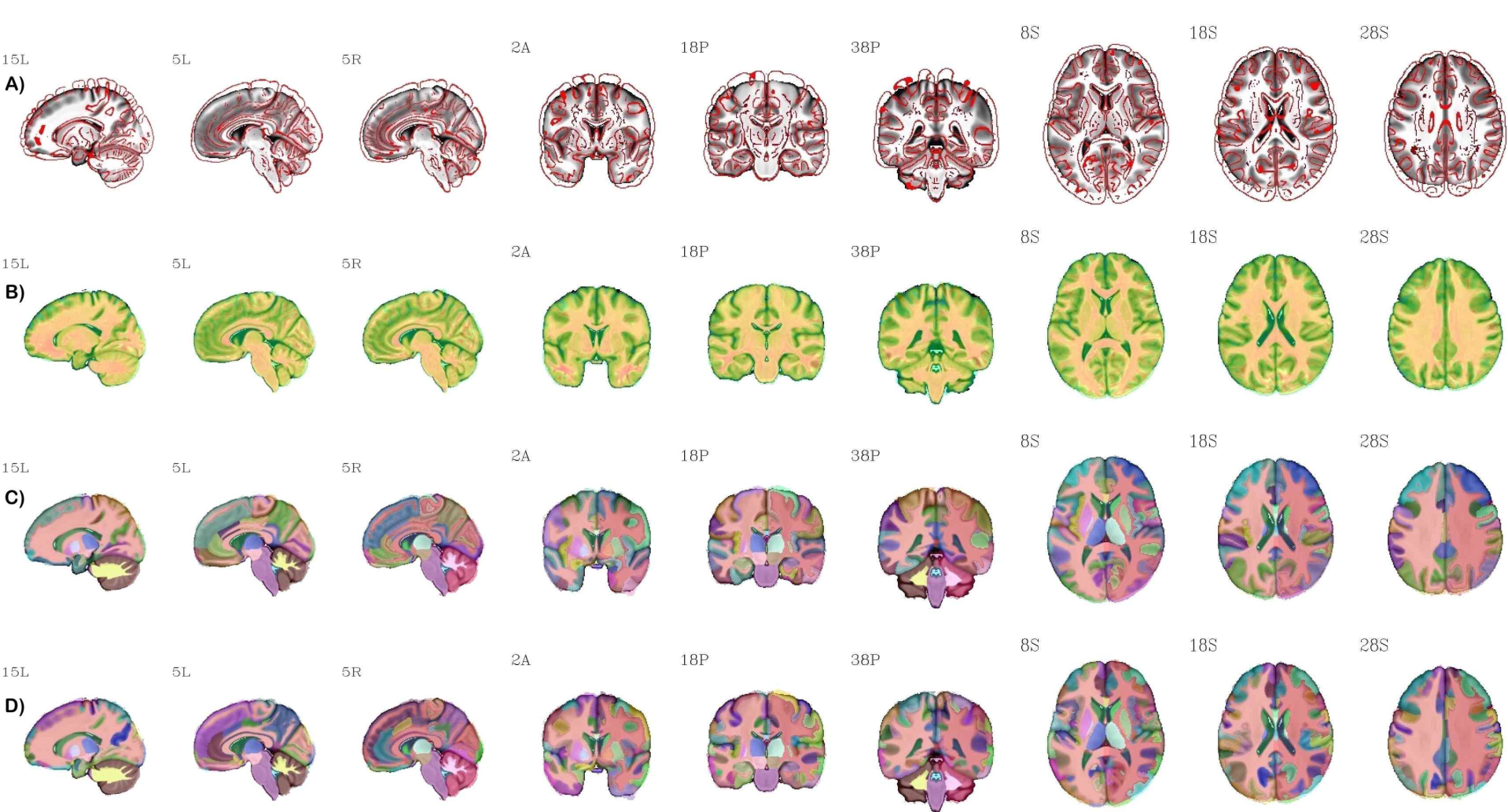
Three sets of sagittal, coronal and axial views of the “population-average” C3 IBT, displayed as underlay in grayscale in each row (A-D). Row A depicts the edge-filtered version of the MNI 2009 nonlinear template as overlay for size comparison. Row B shows the “typical” IBT C3 dataset as a translucent overlay; note the very high degree of structural similarity, as expected. The Indian MPM version of the DK atlas (FreeSurfer’s 2000 atlas) is shown in row C as overlay and Destrieux atlas (FreeSurfer’s 2009 atlas) as overlay in row D.

Figure 4‘s left panel displays the logarithm of the relative volume ratio of each ROI in the IBT MPM atlas (see Eq. (1)), showing how representative the atlas is of each cohort in a region-wise manner. As shown in the figure, most cortical regions have values close to zero, indicating that MPM ROIs in the IBT space provide representative volumes of the native space ROIs for each age group. The largest expansions were observed in the bilateral caudal and rostral middle frontal gyrus, bilateral rostral anterior cingulate, bilateral superior and inferior parietal cortices across the age groups. These are also the regions that show greater mDV (Figure 4‘s right-panel) indicating that greater inter-subject variability could be in part responsible for greater volumetric differences between native-space and MPM volumes. The scatter-plots in Supplementary Information (Figure S7) indicates that there were significant correlations between relative volume ratios and mDV for each age group (*R*-values: 0.24-42 and *p*-values <0.05).

**Figure 4.**
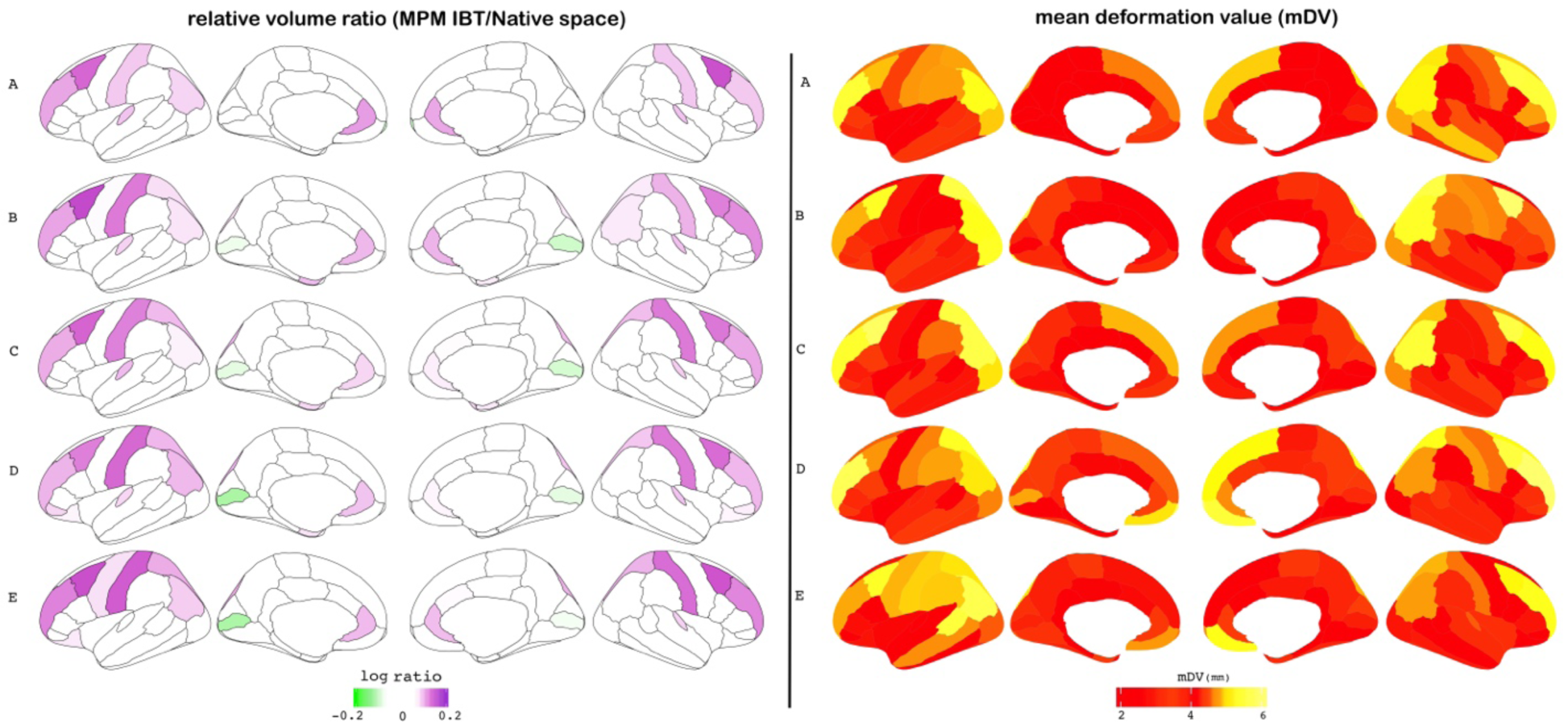
Evaluation of the region-wise similarity of the MPM volumes as measured (left panel) by the relative volume ratio for each ROI via Eq. (1), and (right panel) by mean deformation value (mDV) of each ROI; rows A-E show results for each age-specific group C1-C5, respectively. In the left-panel ROIs with notably different volume fractions are highlighted in purple (increases) and green (decreases), and in the right-panel ROIs with greater inter-subject variability are shown as increasingly yellow.

Figure 5A-E shows the comparison of warp distances from the anatomical (T1w) volumes of the validation cohorts (V1-5) to each of the age-matched IBT “population mean” templates (orange), vs the V1-5 warp distances to the standard MNI template (blue); for more detailed comparison, average warp distances along each of the main volumetric axes are shown separately. In all cases, alignment to an IBT dataset required much less overall displacement on average. Warps to MNI were highly significantly greater (*p<* 0.05, corrected for *N* = 3 × 5 multiple comparisons) along the PA and IS axes in all cases. Along the LR axes, differences were smaller but still significant at the same level for 4/5 cohorts (again, warps to MNI being larger); the C4 cohort showed no significant difference along the LR axis, but overall differences for this group were still large, due to the warps along the other axes.

**Figure 5.**
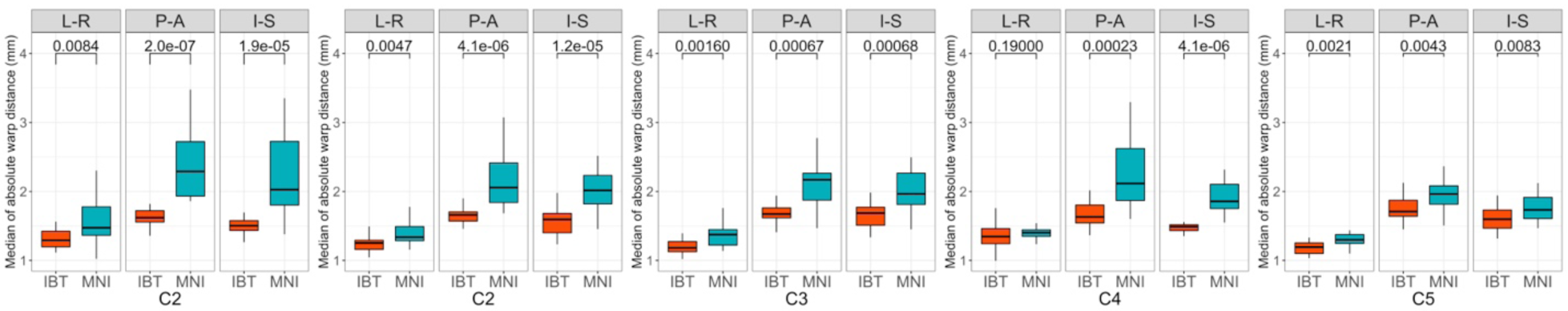
Validation cohort T1w results: A-E) IBT-based results are in orange, and MNI-based results in blue. Wilcoxon’s signed-ranks test was used to compare the distributions; *p*-values are shown at the top of each panel. For each validation group (V1-5), boxplots of the median warp magnitude along each major axis (LR, PA, IS) to a given template are shown in panel A-E. The warp distributions to MNI space are significantly larger along the AP and IS axes in all cases. While the differences tend to be smallest along the LR axis (particularly for C4), warps to MNI are nevertheless significantly larger for 4/5 of the cohorts along this axis, as well.

Finally, we investigated the practical difference when using IBT vs MNI as a template space for fMRI processing, using the validation cohorts. ReHo values were compared between corresponding ROIs in the IBT and MNI spaces, and the paired t-tests of the values showed that each IBT tended to have higher ReHo values throughout most regions of the brain. These results are shown in Figure 6. While some medial and anterior regions showed higher ReHo in the MNI space, the overall greater ReHo values in the IBT space may be the result of slightly improved alignments on average, so that more similar time series are grouped together per ROI.

**Figure 6.**
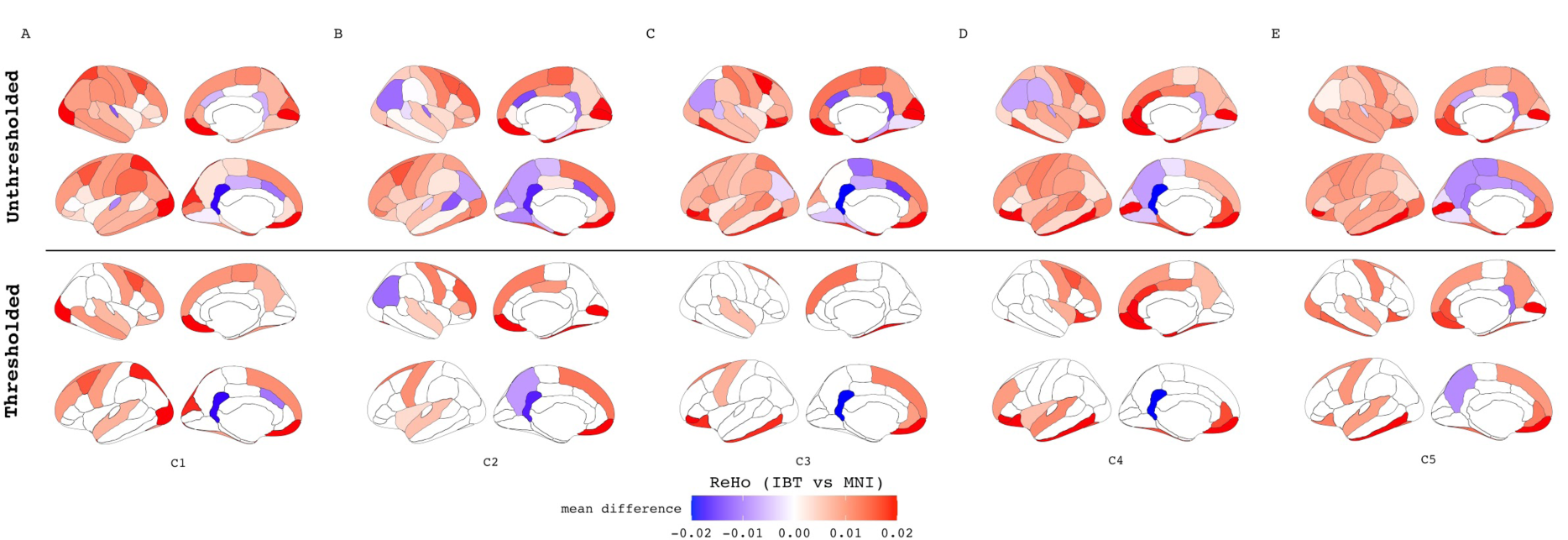
Validation cohort fMRI results. A-E) Comparison of the region-wise ReHo values in the IBT vs MNI space for each validation group C1-C5. The colors indicate the directions and magnitude of the mean difference of ReHo values between IBT vs MNI. The unthresholded results are in top panel and Bonferroni corrected results are in the bottom panel. The warm-red color indicates regions where the ReHo values are greater in IBT and cool-blue colors are those where ReHo values are greater in MNI. The ReHo provides a measure of local FC as index of temporal coherence (Kendall’s coefficient of concordance) of the BOLD time series within a set of a given voxel’s nearest neighbors in an ROI.

## 4 Discussion

We have introduced five new India brain template (IBTs) spaces, spanning an age range from 6-60 years. Additionally, corresponding atlases (IBTAs) from widely used segmentations were also created for each space. These should form useful reference templates and region maps for brain imaging studies involving predominantly Indian populations. Both the creation of age-specific templates and the inclusion of associated atlases make the present study distinct from previous Indian population brain template projects [Rao et al., 2017, Bhalerao et al., 2018, Sivaswamy et al., 2019, Pai et al., 2020]; additionally, we have generated both “population mean” and high-contrast “typical” templates for each age band. The IBT volumes and corresponding atlases are publicly available for download, in standard NIFTI format, and freely usable by the wider neuroimaging community.

The need for age-specific templates in particular has been recognized across different populations [Fonov et al., 2011, Wilke et al., 2002, Yoon et al., 2009]; however, Indian versions of age-specific brain templates have not been available to date. While adult brain templates may still provide reasonably accurate anatomical priors for normalizing lower resolution smoothed functional data, they may not be appropriate for high resolution structural and functional data [Wilke et al., 2002]. For example, Yoon et al. [2009] examined the “template effect” in a pediatric population and noted significantly greater amount of deformation required for nonlinear normalization to the MNI152 adult template than compared to an age-appropriate template (2.2 vs. 1.7 mm). Further, the authors also noted significant differences in both volume-based and surface-based morphological features between data warped to pediatric and adult brain templates. Such discrepancies are also reported in aging studies, where use of young-adult template (such as the MNI) for older adults can result in biases such as regional distortion and systematic over-expansion of older brains [Buckner et al., 2004]. Age-appropriate template for older adults have also been shown to provide more accurate tissue segmentation for structural imaging [Fillmore et al., 2015] and more focused activation patterns with improvement in sensitivity for fMRI group analyses [Huang et al., 2010].

In addition to age, consideration should also be given to the ethnic or population-specific differences [Lee et al., 2005, Tang et al., 2010, Rao et al., 2017], when choosing the appropriate brain template. As expected, there are noticeable structural differences when comparing the new IBTs with existing, popular standard templates (such as the MNI), which have been made from very different subject populations. Overall, registration to the IBTs from the Indian population validation groups required much less deformation of the input datasets and resulted in more accurate stereotactic standardization and anatomical localization. The relative differences in warping along the major axes of the brain were shown here using validation groups from the local population. The differences in warping magnitudes varied both by axis and by the age of subjects. Thus, the structural differences in templates are not trivial, i.e., just scaling, but instead reflect shape variations that are likely to significantly affect the overall goodness-of-fit and anatomical alignment across a group study.

Such aspects were highlighted in the differences of outcomes in fMRI processing when using IBT vs MNI templates: the IBT-based output tended to have higher ReHo values among ROI pairs. The latter fact in particular suggests that the IBTs provided better function-to-anatomical alignment across groups, so that voxel with functionally similar time series tended to be grouped together more preferentially. One might expect this to be a relatively small effect, because alignment to the MNI templates still appears generally reasonable; one would expect the overlap pattern differences to be occurring fractionally within ROIs and predominantly at boundaries. Indeed, the FC differences were relatively small, but with a noticeable trend toward higher values in the IBT-based datasets.

It is important to emphasize that these structural differences are only with regards to morphology; they do not relate to functional or behavioral outcomes, nor to intelligence, etc. The purpose and goal of population-specific templates is for the practical consideration of maximizing the matching of structures across a group during an alignment step of processing, as well as to better match functional regions to structures. These are geometric and signal-to-noise considerations, which are important in brain studies (as demonstrated here), but which are unrelated to the brain behavior itself.

The wide variety of brain structural patterns in any group, even in an apparently homogeneous one, is also worth commenting on. This inherent variability affects both the creation and utilization of brain templates [Yang et al., 2020]. In any population brain structures can vary to the degree of having different numbers of sulci in the same region (e.g., [Thompson et al., 1996] and *op cit*); this is true even in a group of controls who are highly localized, genetically related, similar age and background, etc. Thus, there is a minimum and nontrivial degree of variability in alignment that one can reasonably expect both when combining multiple subjects to generate a template, as well as in the overlap of anatomical structures when applying the template. Indeed, the Indian population (currently over 1.3 billion people) is spread across a wide range of geographies with diversity in linguistic-ethnic compositions as well as extensive genetic admixtures [Basu et al., 2016]. In this study, the final mean template for each cohort contained variability. However, this was relatively low compared to the mean dataset values, and the final mean template contained a large amount of clearly defined structure. Moreover, the fractional overlap of ROIs when generating the maximum probability map atlases showed a high degree of agreement across the group through most of the brain.

The variability present in the template generation is also observable in the atlases. The inter-subject variability (as measured by the mean deformation values for various regions during non-linear registration to age and population-specific template) also correlated positively with the expansion of MPM volumes, in all age groups (see Supplementary Figure S7). While the final MPM atlases indicate the most frequent positions of each brain region in a given cohort, we also provide the probability density maps for each ROI in the atlas (see supplementary Figure S8 for example), which can be of additional use in ROI-based analyses.

While spatial normalization to IBT offers distinct advantages in terms of spatial accuracy and detection power, it may still be desirable to have the results from any particular analysis also reported in another space. For example, for comparisons with previously published studies, one might want to compare the locations of a finding with those reported in MNI, Talairach or Korean template coordinate spaces. Therefore, a nonlinear coordinate transformation mapping between IBT and the common MNI space has also been calculated, and a similar coordinate warp between *any* coordinate frames can be calculated easily.

There are several methodological strengths and limitations related to the current study that should be noted. We used combined state-of-the-art linear and non-linear averaging techniques using AFNI’s completely automated pipeline “make_template_dask.py”, which uses the Dask python parallelization to efficiently make a template from a large group of subjects. We addressed several specific challenges involved in the template creation, such as intensity normalization from different scanners, scaling, resizing of the overall brain size to be representative of the cohort at each iteration, and anisotropic smoothing with preservation of edges. While the overall sample size of the study was relatively large, the late childhood and the late adulthood templates had relative modest sample sizes. Therefore, it will be of benefit for the constructed templates to continue to be updated with larger sample sizes as we collect more MRI datasets. Future work should also expand the templates for ages *<* 6 yr and *>* 60 yr. We will also expand this work to include development of a cortical surface atlas, which may allow for a registration procedure involving alignment of highly variable cortical folding patterns.

## 5 Conclusions

In conclusion, the present work demonstrates the appropriateness of using age and population-specific templates as reference targets for spatial normalization of structural and functional neuroimaging data. This database of age-specific IBTs and IBTAs is made freely available to the wider neuroimaging community of researchers and clinicians worldwide. We hope that these tools will facilitate research into neurological understand in general and into the functional and morphometric changes that occur over life-course in Indian population in particular.

## Data Availability Statement

The Indian brain templates (IBTs) and atlases (IBTAs) developed in this study are openly available for use in AFNI. Instructions for downloading the datasets are available at https://hollabharath.github.io/IndiaBrainTemplates. The installer script is also available from Zenodo at https://doi.org/10.5281/zenodo.3817045.

## Author Contributions

VB, RDB, BH, PAT and DRG conceptualized and designed the study. VB, RDB, PP, GV, UMM, JS, MK, KK, AC and DB contributed data to the study. BH, PAT, NV and DPO curated the data. BH and PAT conducted data quality assessments. BH, PAT, DRG and JAL conducted the computations required for template construction. GV and NPR contributed data for the validation experiments. BH and PAT conducted the validation experiments. BH and PAT took the lead in writing the manuscript. DRG, GJB, RDB, RWC and VB contributed to the interpretation of the findings and edited the manuscript for important intellectual content. All authors discussed the results and contributed to the final manuscript.

## Acknowledgments

This work was partially supported by c-VEDA (Consortium on Vulnerability to Externalizing Disorders and Addictions) ICMR (India)/MRC (UK) (grant ICMR/MRC-UK/3/M/2015-NCD-I) to VB and GS. Wellcome Trust/DBT India Alliance Fellowship Grants to BH (Award: IA/RTF/14/1/1002), and UMM (Award: IA/E/12/1/500755), DST Research Grant SR/CSI/44/2008(5) to RDB and DBT Research Grant BT/PR14315/MED/30/474/2010 to PKP. GV acknowledges the support of the SwarnaJayanti Fellowship by the Department of Science and Technology, Government of India (DST/SJF/LSA-02/2014–15). GS was supported by the Horizon 2020-funded ERC Advanced Grant ‘STRATIFY’ (brain network-based stratification of reinforcement-related disorders; 695313), ERANID (understanding the interplay between cultural, biological and subjective factors in drug use pathways; PR-ST-0416-10004), BRIDGET (JPND brain imaging, cognition, dementia and next generation GEnomics; MR/N027558/1), the Human Brain Project (SGA 2, 785907, and SGA 3, 945539), the National Institute of Health (NIH) (R01DA049238, A decentralized macro and micro gene-by-environment interaction analysis of substance use behavior and its brain biomarkers). DRG, JAL, PAT and RWC were supported by the NIMH and NINDS Intramural Research Programs (ZICMH002888) of the NIH (HHS, USA). This work utilized the computational resources of the NIH HPC Biowulf cluster (http://hpc.nih.gov).

## Supplementary Information

This section provides supplementary figures and codes to the material in the main text.

**The contrast-to-noise ratio (CNR)** between GM and WM for successive template creation steps (Figure S1) was calculated as

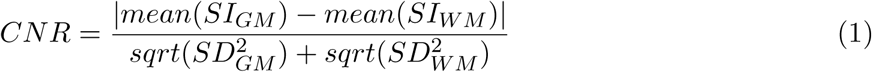

where SI_GM_ / SI_wM_ and SD_GM_/SD_WM_ are mean signal intensity within the gray and white matter respectively, and the corresponding standard deviations.

**Figure S1:**
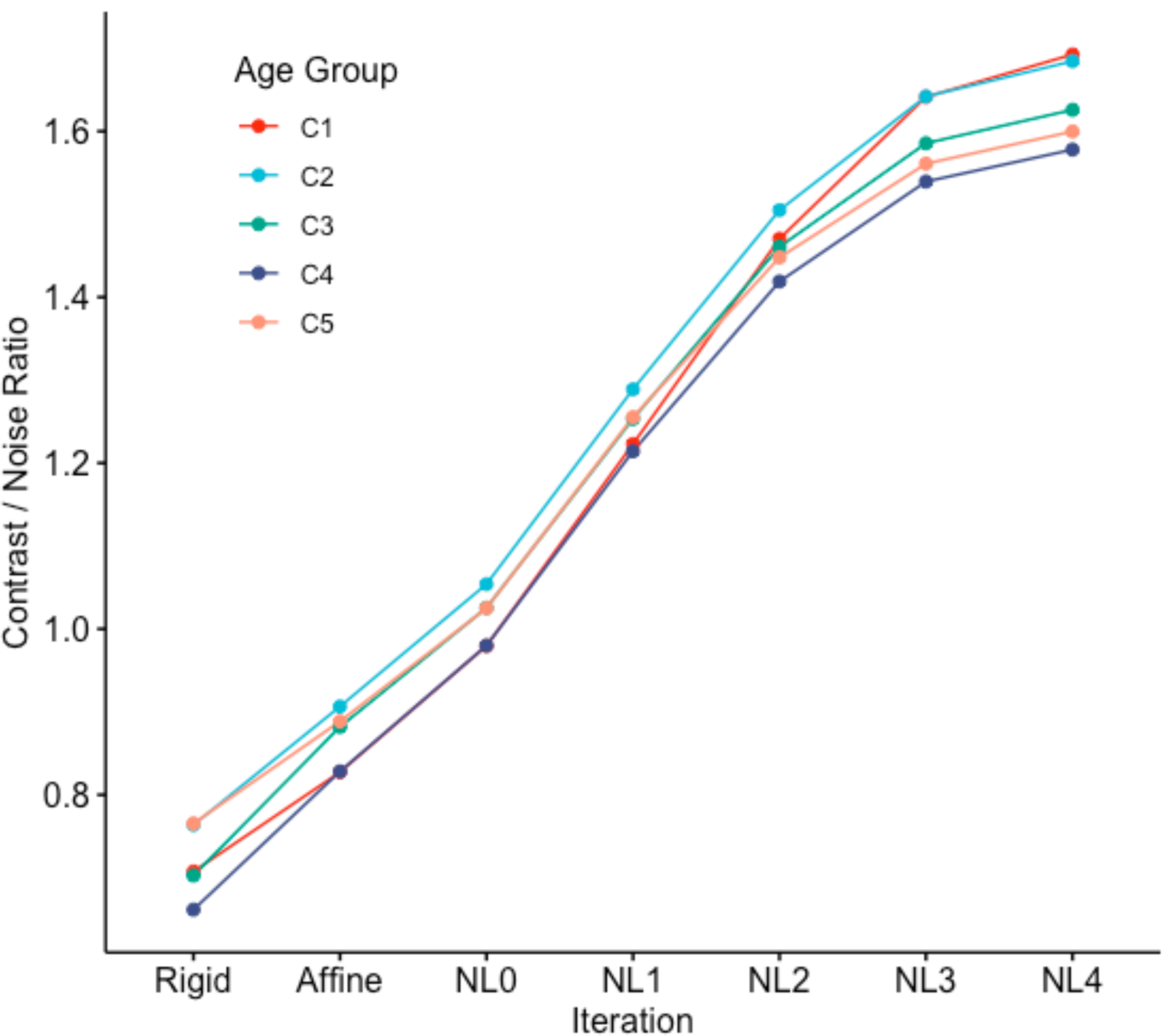
The CNR between GM and WM improves consistently across the successive template creation stages in all the template age-groups C1-C5.

**Figure S2:**
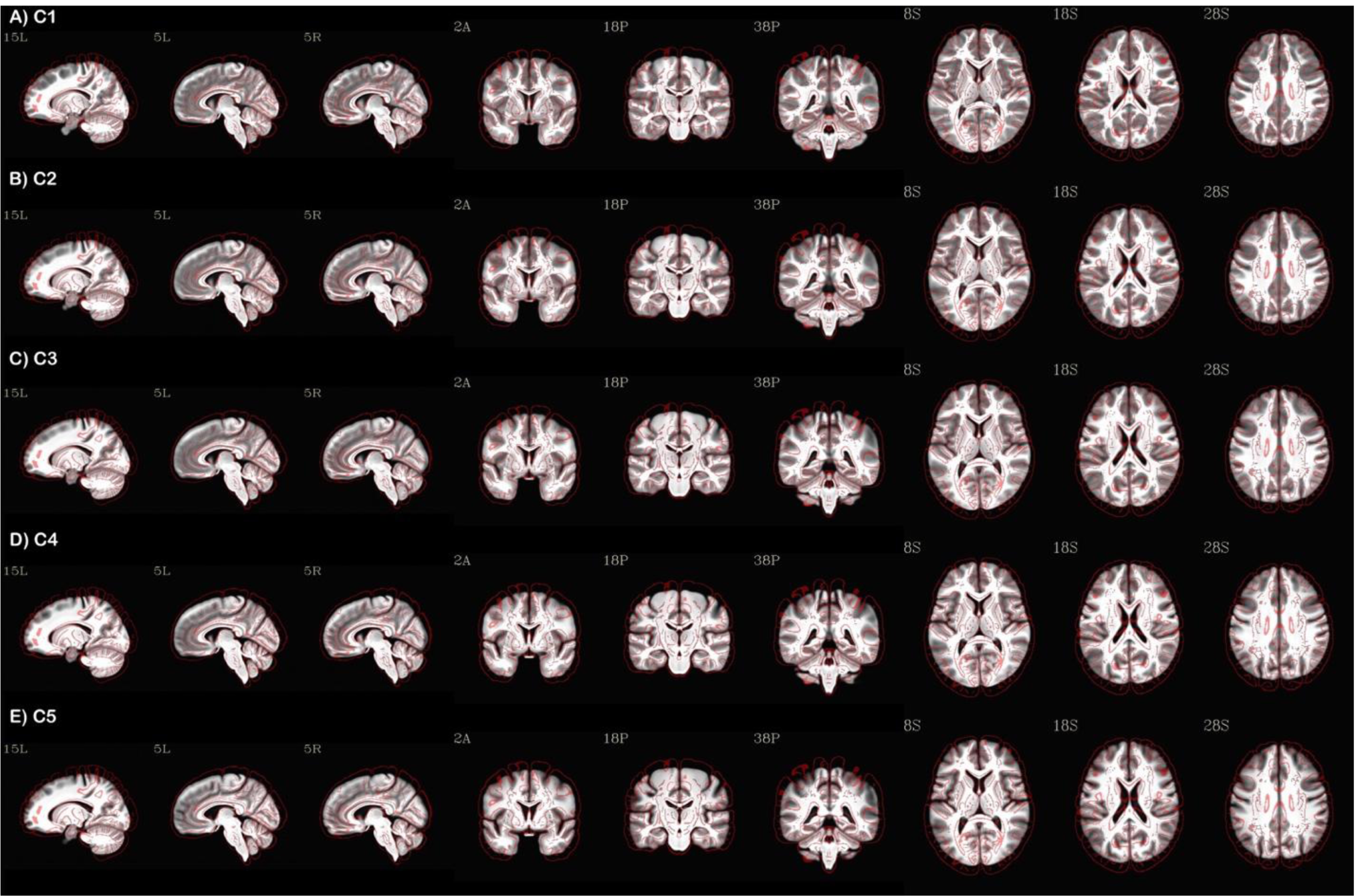
The five IBTs (C1-5) with three sets of sagittal, coronal and axial view displayed as underlay in grayscale and edge-filtered version of the MNI 2009 non-linear template mask as overlay for size comparison. High tissue contrast and detail are evident in each case.

**Figure S3:**
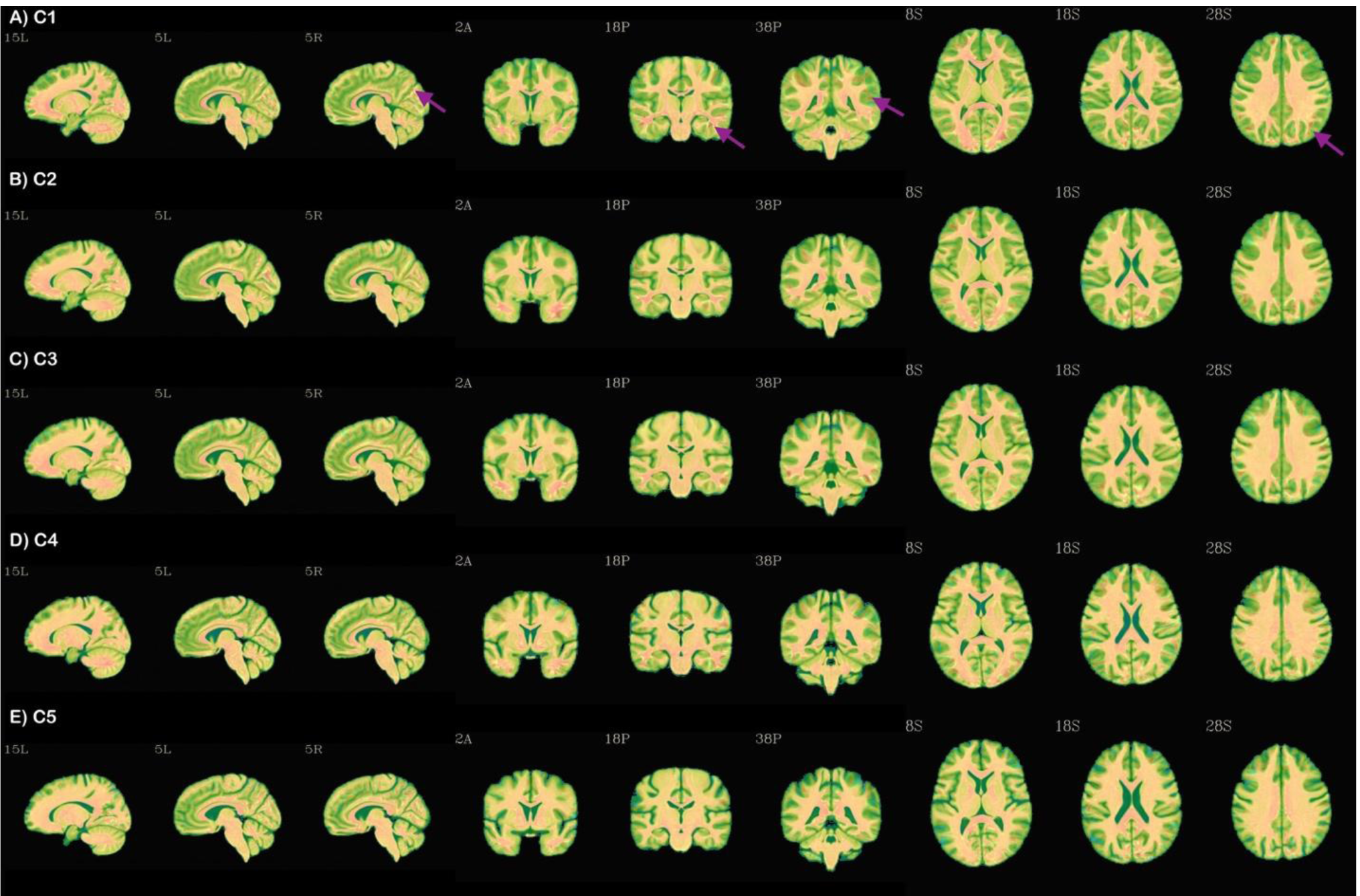
The five population-average IBTs (C1-5) with three sets of sagittal, coronal and axial view displayed as underlay in grayscale and the respective typical subject for each IBT version as the overlay. Arrow points to example regions in C1 age-band regions where the typical version provides greater details than the underlying population-average version.

**Figure S4:**
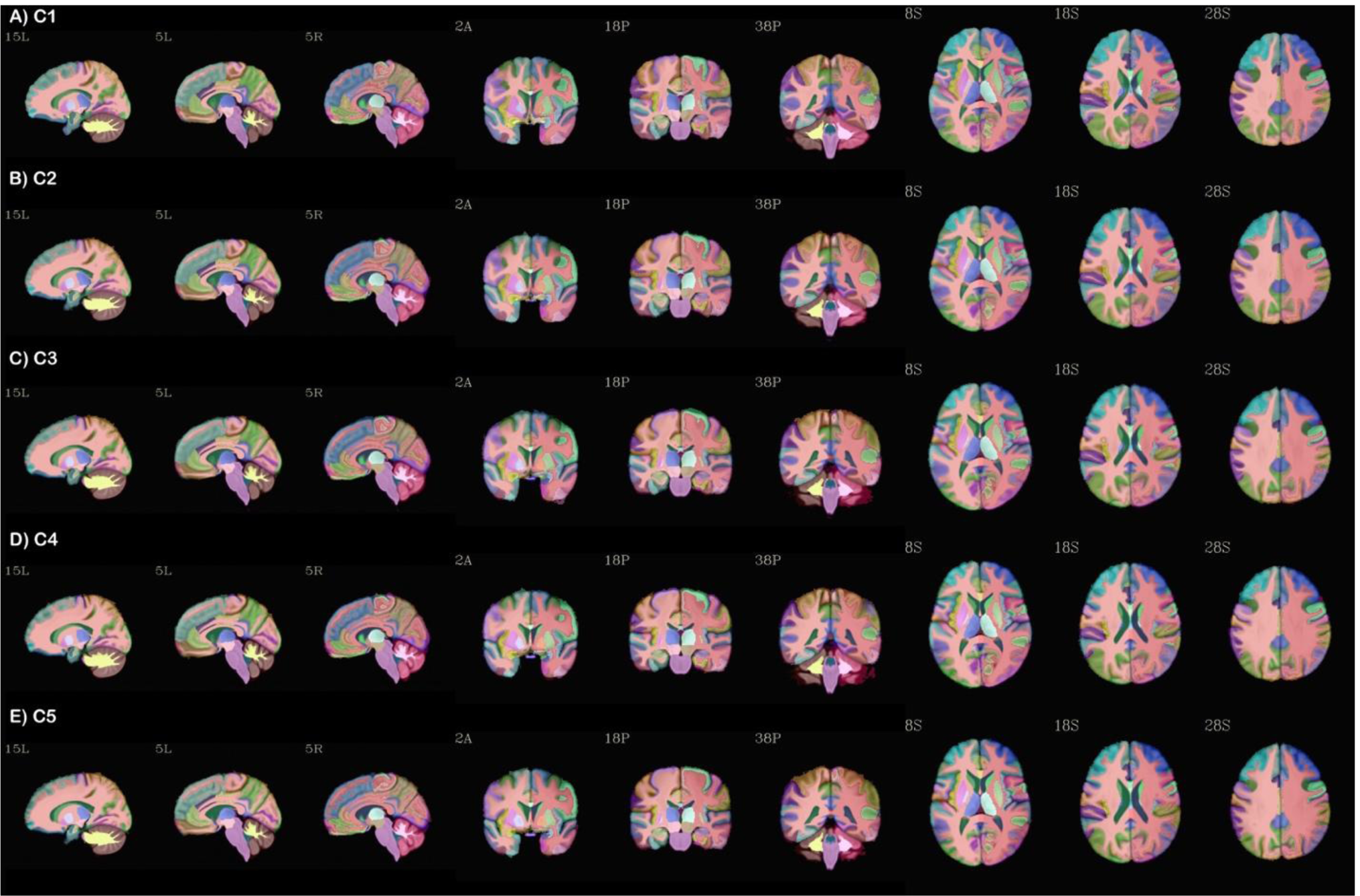
The five IBTs (C1-5) with three sets of sagittal, coronal and axial view displayed as underlay in grayscale and the respective Indian maximum probability map version of the DK atlas (FreeSurfer’s 2000 Atlas) as overlay in AFNI’s “ROI_i256” color scale.

**Figure S5:**
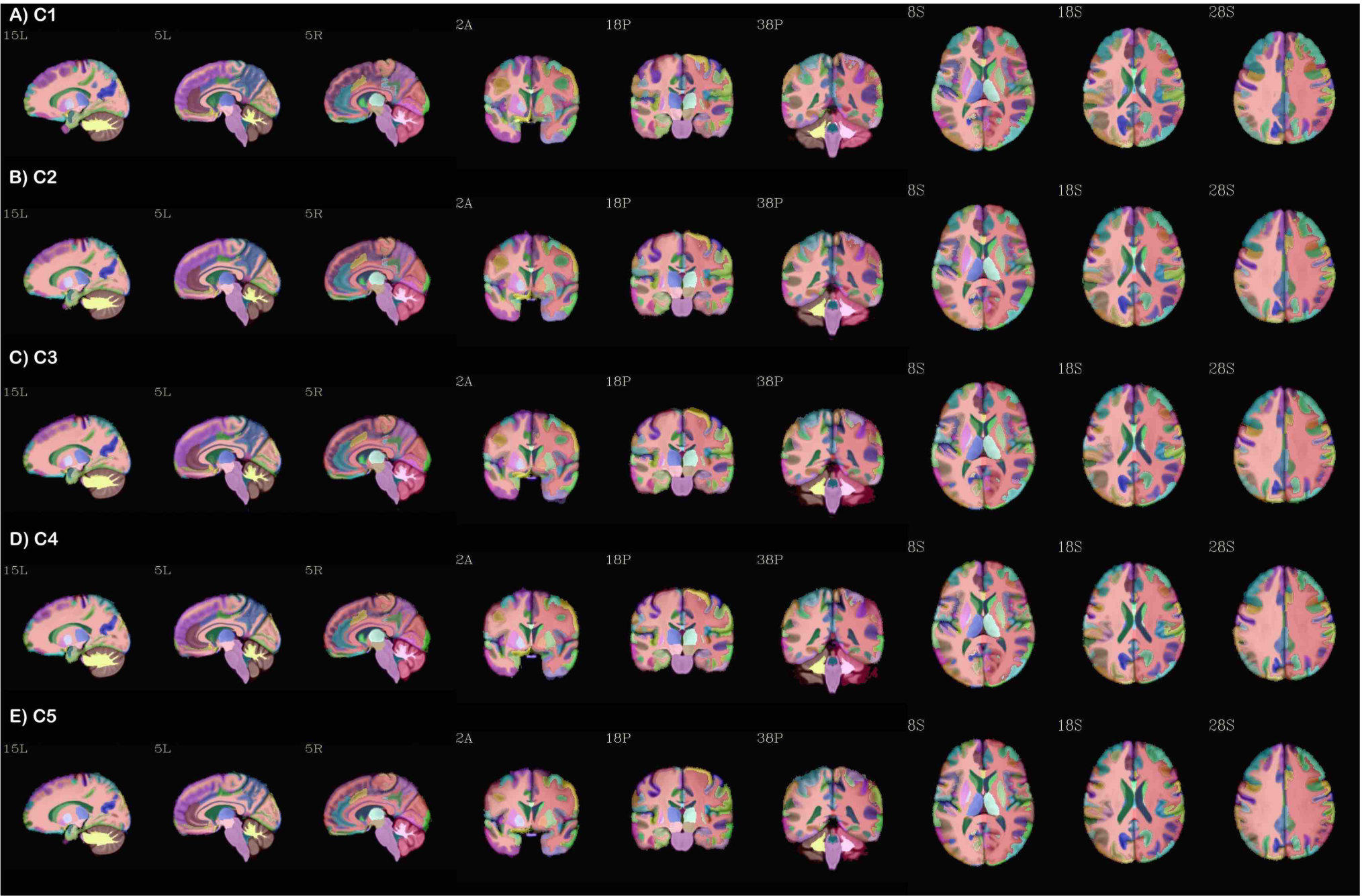
The five IBTs (C1-5) with three sets of sagittal, coronal and axial view displayed as underlay in grayscale and the respective Indian maximum probability map version of the Destrieux atlas (FreeSurfer’s 2009 Atlas) as overlay in AFNI’s “ROI_i256” color scale.

**Figure S6:**
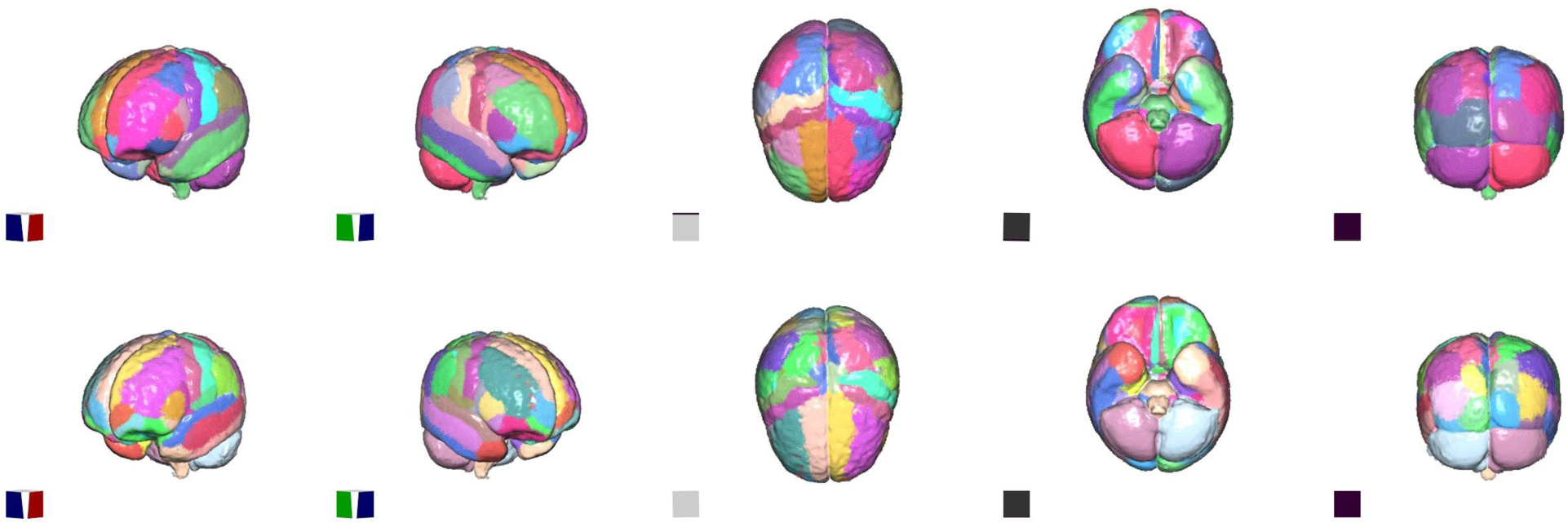
3D surface view of the brain atlases for the C1-IBT age band. The top row shows the maximum probability map (MPM) version of the DK atlas (FreeSurfer’s 2000 Atlas) and the bottom row shows MPM version of the Destrieux atlas (FreeSurfer’s 2009 Atlas) for the C1 age band.

**Figure S7:**
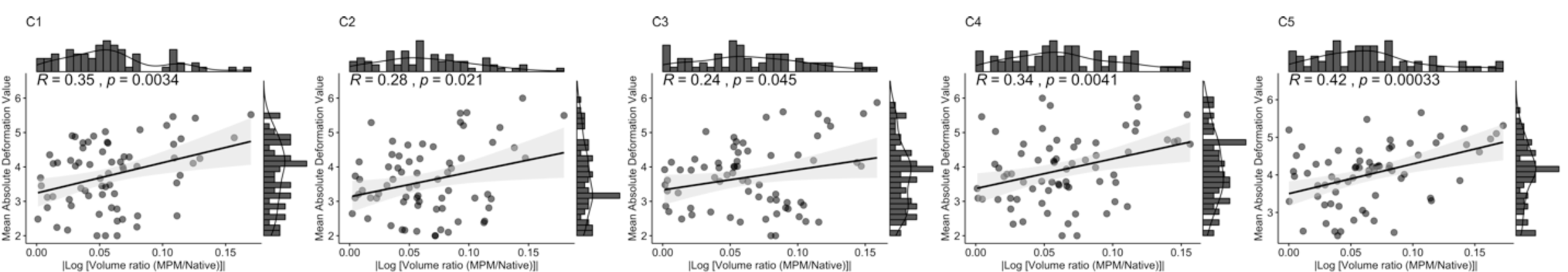
Scatterplot with marginal densigram for pairwise correlations between absolute values of logarithm of the relative volume ratios and mean absolute deformation value across all the regions in the maximum probability map (MPM) version of the DK atlas (FreeSurfer’s 2000 Atlas) at each age-group C1-C5.

**Figure S8:**
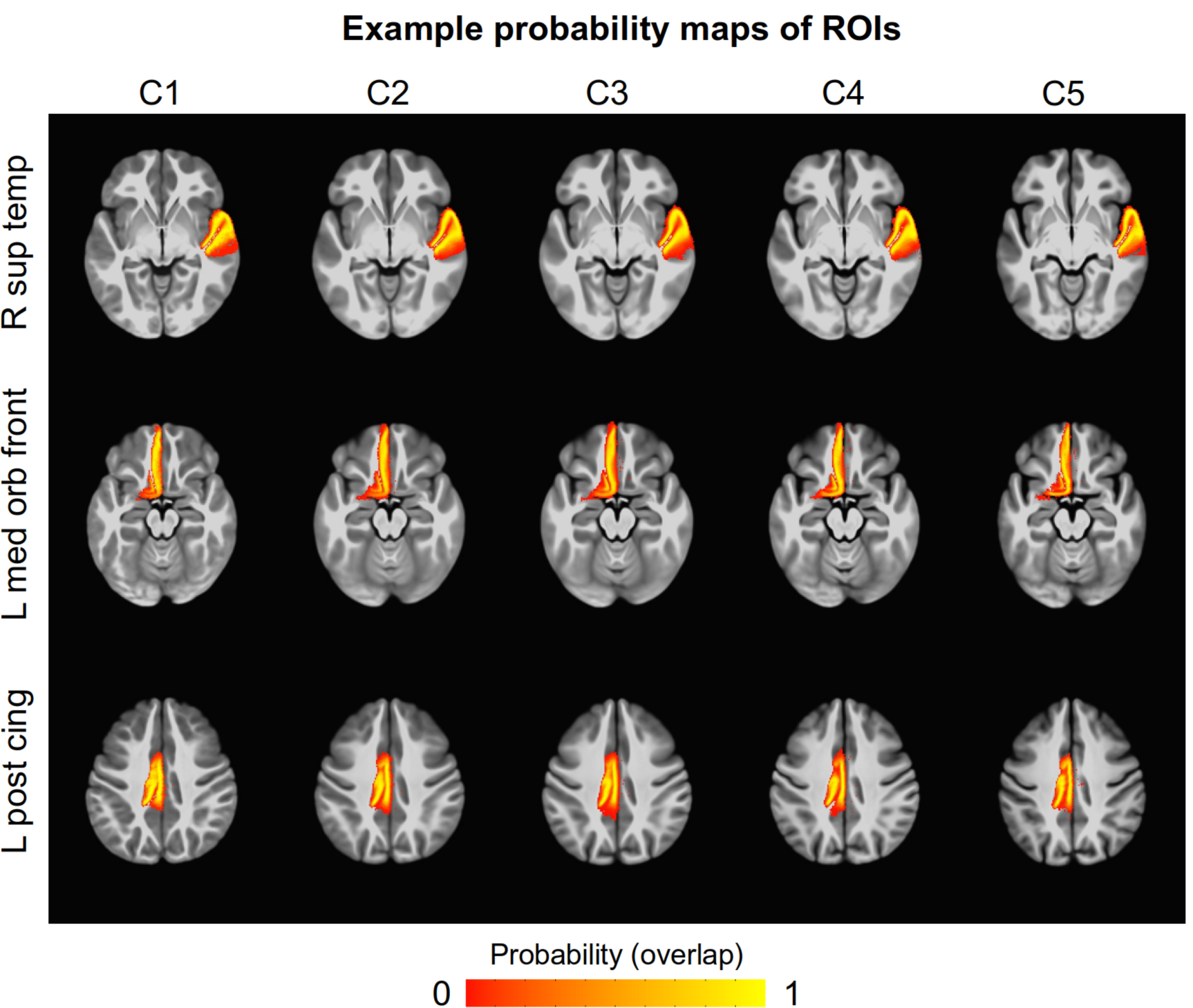
Axial views for three example region of interest from MPM-2000 IBT atlas for all the age groups. The top row shows probability map for right superior temporal gyrus, middle row shows left medial orbital frontal gyrus and the bottom row shows left posterior cingulate gyrus. The color intensity reflects probability density estimates (ranging from 0 to 1)

## Supplementary Information

Example afni_proc.py command for comparing validation tests.

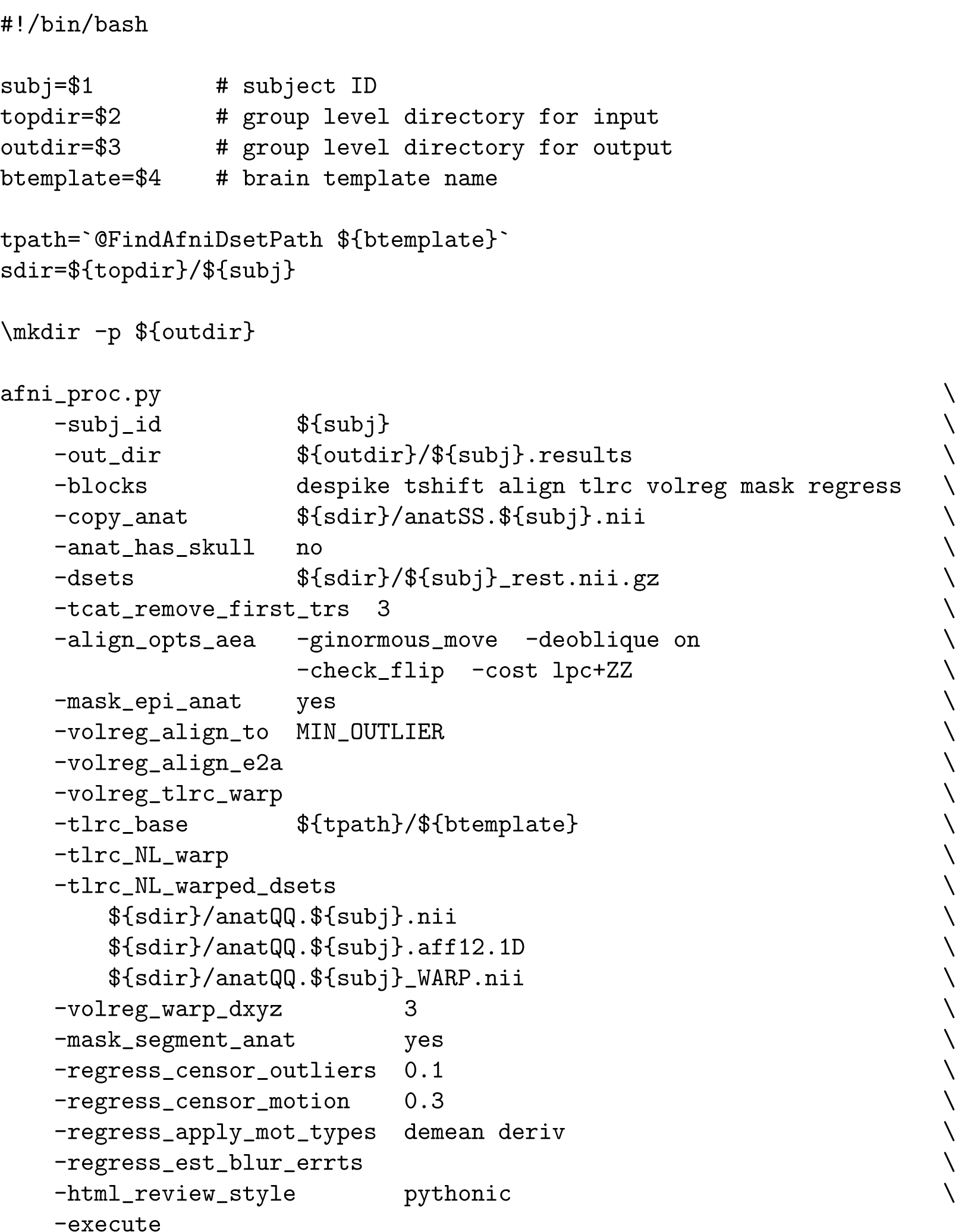

## Footnotes

1 FreeSurfer distinguishes between cortical parcellations and subcortical segmentations; here, we use “parcellation” generically to refer to final map of all ROIs.

## References

Analabha Basu, Neeta Sarkar-Roy, and Partha P. Majumder. Genomic reconstruction of the history of extant populations of india reveals five distinct ancestral components and a complex structure. Proceedings of the National Academy of Sciences, 113(6):1594–1599, January 2016. doi: 10.1073/pnas.1513197113. URL https://doi.org/10.1073/pnas.1513197113.

G. V. Bhalerao, R. Parlikar, R. Agrawal, V. Shivakumar, S. V. Kalmady, N. P. Rao, S. M. Agarwal, J. C. Narayanaswamy, Y. C. J. Reddy, and G. Venkatasubramanian. Construction of population-specific indian mri brain template: Morphometric comparison with chinese and caucasian templates. Asian J Psychiatr, 35:93–100, 2018. ISSN 1876-2026 (Electronic) 1876-2018 (Linking). doi: 10.1016/j.ajp.2018.05.014. URL https://www.ncbi.nlm.nih.gov/pubmed/29843077.

Randy L. Buckner, Denise Head, Jamie Parker, Anthony F. Fotenos, Daniel Marcus, John C. Morris, and Abraham Z. Snyder. A unified approach for morphometric and functional data analysis in young, old, and demented adults using automated atlas-based head size normalization: reliability and validation against manual measurement of total intracranial volume. NeuroImage, 23(2):724–738, October 2004. doi: 10.1016/j.neuroimage.2004.06.018. URL https://doi.org/10.1016/j.neuroimage.2004.06.018.

R. W. Cox. Afni: software for analysis and visualization of functional magnetic resonance neuroimages. Comput Biomed Res, 29(3):162–73, 1996. ISSN 0010-4809 (Print) 0010-4809 (Linking). doi: 10.1006/cbmr.1996.0014. URL https://www.ncbi.nlm.nih.gov/pubmed/8812068.

RW Cox and DR Glen. Nonlinear warping in afni. In Poster presented at the 19th Annual Meeting of the Organization for Human Brain Mapping, 2013.

Dask Development Team.x Dask: Library for dynamic task scheduling, 2016. URL https://dask.org.

R. S. Desikan, F. Segonne, B. Fischl, B. T. Quinn, B. C. Dickerson, D. Blacker, R. L. Buckner, A.M. Dale, R. P. Maguire, B. T. Hyman, M. S. Albert, and R. J. Killiany. An automated labeling system for subdividing the human cerebral cortex on mri scans into gyral based regions of interest. Neuroimage, 31(3):968–80, 2006. ISSN 1053-8119 (Print) 1053-8119 (Linking). doi: 10.1016/j.neuroimage.2006.01.021. URL https://www.ncbi.nlm.nih.gov/pubmed/16530430.

Christophe Destrieux, Bruce Fischl, Anders Dale, and Eric Halgren. Automatic parcellation of human cortical gyri and sulci using standard anatomical nomenclature. Neuroimage, 53(1):1–15, 2010. ISSN 1053-8119.

Alan C Evans, D Louis Collins, SR Mills, ED Brown, RL Kelly, and Terry M Peters. 3d statistical neuroanatomical models from 305 mri volumes. In 1993 IEEE conference record nuclear science symposium and medical imaging conference, pages 1813–1817. IEEE, 1993.

Paul T. Fillmore, Michelle C. Phillips-Meek, and John E. Richards. Age-specific MRI brain and head templates for healthy adults from 20 through 89 years of age. Frontiers in Aging Neuroscience, 7, April 2015. doi: 10.3389/fnagi.2015.00044. URL https://doi.org/10.3389/fnagi.2015.00044.

Fischl. Freesurfer. Neuroimage, 62(2):774–81, 2012. ISSN 1095-9572 (Electronic) 1053-8119 (Linking). doi: 10.1016/j.neuroimage.2012.01.021. URL https://www.ncbi.nlm.nih.gov/pubmed/22248573.

V. Fonov, A. C. Evans, K. Botteron, C. R. Almli, R. C. McKinstry, D. L. Collins, and Group Brain Development Cooperative. Unbiased average age-appropriate atlases for pediatric studies. Neuroimage, 54(1):313–27, 2011. ISSN 1095-9572 (Electronic) 1053-8119 (Linking). doi: 10.1016/j.neuroimage.2010.07.033. URL https://www.ncbi.nlm.nih.gov/pubmed/20656036.

Chih-Mao Huang, Shwu-Hua Lee, Ing-Tsung Hsiao, Wan-Chun Kuan, Yau-Yau Wai, Han-Jung Ko, Yung-Liang Wan, Yuan-Yu Hsu, and Ho-Ling Liu. Study-specific EPI template improves group analysis in functional MRI of young and older adults. Journal of Neuroscience Methods, 189(2): 257–266, June 2010. doi: 10.1016/j.jneumeth.2010.03.021. URL https://doi.org/10.1016/j.jneumeth.2010.03.021.

G. Kendall and B. Babington Smith. The problem of m rankings. Ann. Math. Statist., 10 (3):275–287, 09 1939. doi: 10.1214/aoms/1177732186. URL https://doi.org/10.1214/aoms/1177732186.

Jae Sung Lee, Dong Soo Lee, Jinsu Kim, Yu Kyeong Kim, Eunjoo Kang, Hyejin Kang, Keon Wook Kang, Jong Min Lee, Jae-Jin Kim, and Hae-Jeong Park. Development of korean standard brain templates. Journal of Korean medical science, 20(3):483–488, 2005. ISSN 1011-8934.

X. Li, P. S. Morgan, J. Ashburner, J. Smith, and C. Rorden. The first step for neuroimaging data analysis: Dicom to nifti conversion. J Neurosci Methods, 264:47–56, 2016. ISSN 1872-678X (Electronic) 0165-0270 (Linking). doi: 10.1016/j.jneumeth.2016.03.001. URL https://www.ncbi.nlm.nih.gov/pubmed/26945974.

John Mazziotta, Arthur Toga, Alan Evans, Peter Fox, Jack Lancaster, Karl Zilles, Roger Woods, Tomas Paus, Gregory Simpson, Bruce Pike, et al. A four-dimensional probabilistic atlas of the human brain. Journal of the American Medical Informatics Association, 8(5):401–430, 2001a.

John Mazziotta, Arthur Toga, Alan Evans, Peter Fox, Jack Lancaster, Karl Zilles, Roger Woods, Tomas Paus, Gregory Simpson, Bruce Pike, et al. A probabilistic atlas and reference system for the human brain: International consortium for brain mapping (icbm). Philosophical Transactions of the Royal Society of London. Series B: Biological Sciences, 356(1412):1293–1322, 2001b.

Praful P. Pai, Pravat K. Mandal, Khushboo Punjabi, Deepika Shukla, Anshika Goel, Shallu Joon, Saurav Roy, Kanika Sandal, Ritwick Mishra, and Ritu Lahoti. BRAHMA: Population specific t1, t2, and FLAIR weighted brain templates and their impact in structural and functional imaging studies. Magnetic Resonance Imaging, 70:5–21, July 2020. doi: 10.1016/j.mri.2019.12.009. URL https://doi.org/10.1016/j.mri.2019.12.009.

P. Rao, H. Jeelani, R. Achalia, G. Achalia, A. Jacob, R. D. Bharath, S. Varambally, G. Venkatasubramanian, and K. Yalavarthy P. Population differences in brain morphology: Need for population specific brain template. Psychiatry Res Neuroimaging, 265:1–8, 2017. ISSN 1872-7506 (Electronic) 0925-4927 (Linking). doi: 10.1016/j.pscychresns.2017.03.018. URL https://www.ncbi.nlm.nih.gov/pubmed/28478339.

Z. S. Saad, D. R. Glen, G. Chen, M. S. Beauchamp, R. Desai, and R. W. Cox. A new method for improving functional-to-structural mri alignment using local pearson correlation. Neuroimage, 44 (3):839–48, 2009. ISSN 1095-9572 (Electronic) 1053-8119 (Linking). doi: 10.1016/j.neuroimage.2008.09.037. URL https://www.ncbi.nlm.nih.gov/pubmed/18976717.

Eesha Sharma, Nilakshi Vaidya, Udita Iyengar, Yuning Zhang, Bharath Holla, Meera Purushottam, Amit Chakrabarti, Gwen Sascha Fernandes, Jon Heron, Matthew Hickman, Sylvane Desrivieres, Kamakshi Kartik, Preeti Jacob, Madhavi Rangaswamy, Rose Dawn Bharath, Gareth Barker, Dimitri Papadopoulos Orfanos, Chirag Ahuja, Pratima Murthy, Sanjeev Jain, Mathew Varghese, Deepak Jayarajan, Keshav Kumar, Kandavel Thennarasu, Debashish Basu, B. N. Subodh, Rebecca Kuriyan, Sunita Simon Kurpad, Kumaran Kalyanram, Ghattu Krishnaveni, Murali Krishna, Rajkumar Lenin Singh, L. Roshan Singh, Kartik Kalyanram, Mireille Toledano, Gunter Schumann, Vivek Benegal, and The cVEDA Consortium. Consortium on vulnerability to externalizing disorders and addictions (cveda): A developmental cohort study protocol. BMC Psychiatry, 20(1):2, 2020. ISSN 1471-244X. doi: 10.1186/s12888-019-2373-3. URL https://doi.org/10.1186/s12888-019-2373-3.

J. Sivaswamy, A. J. Thottupattu, R. Mehta, R. Sheelakumari, and C. Kesavadas. Construction of indian human brain atlas. Neurol India, 67(1):229–234, 2019. ISSN 0028-3886 (Print) 0028-3886 (Linking). doi: 10.4103/0028-3886.253639. URL https://www.ncbi.nlm.nih.gov/pubmed/30860125.

Jean Talairach and Pierre Tournoux. Co-planar stereotaxic atlas of the human brain-3-dimensional proportional system. An approach to cerebral imaging, 1988.

Y. Tang, C. Hojatkashani, I. D. Dinov, B. Sun, L. Fan, X. Lin, H. Qi, X. Hua, S. Liu, and A. W. Toga. The construction of a chinese mri brain atlas: a morphometric comparison study between chinese and caucasian cohorts. Neuroimage, 51(1):33–41, 2010. ISSN 1095-9572 (Electronic) 1053-8119 (Linking). doi: 10.1016/j.neuroimage.2010.01.111. URL https://www.ncbi.nlm.nih.gov/pubmed/20152910.

Paul A Taylor and Ziad S Saad. Fatcat:(an efficient) functional and tractographic connectivity analysis toolbox. Brain connectivity, 3(5):523–535, 2013.

P. M. Thompson, C. Schwartz, R. T. Lin, A. A. Khan, and A. W. Toga. Three-dimensional statistical analysis of sulcal variability in the human brain. J Neurosci, 16(13):4261–74, 1996. ISSN 0270-6474 (Print) 0270-6474 (Linking). URL https://www.ncbi.nlm.nih.gov/pubmed/8753887.

M. Wilke, V. J. Schmithorst, and S. K. Holland. Assessment of spatial normalization of whole-brain magnetic resonance images in children. Hum Brain Mapp, 17(1):48–60, 2002. ISSN 1065-9471 (Print) 1065-9471 (Linking). doi: 10.1002/hbm.10053. URL https://www.ncbi.nlm.nih.gov/pubmed/12203688.

Guoyuan Yang, Sizhong Zhou, Jelena Bozek, Hao-Ming Dong, Meizhen Han, Xi-Nian Zuo, Hesheng Liu, and Jia-Hong Gao. Sample sizes and population differences in brain template construction. NeuroImage, 206:116318, February 2020. doi: 10.1016/j.neuroimage.2019.116318. URL https://doi.org/10.1016/j.neuroimage.2019.116318.

U. Yoon, V. S. Fonov, D. Perusse, A. C. Evans, and Group Brain Development Cooperative. The effect of template choice on morphometric analysis of pediatric brain data. Neuroimage, 45(3): 769–77, 2009. ISSN 1095-9572 (Electronic) 1053-8119 (Linking). doi: 10.1016/j.neuroimage.2008.12.046. URL https://www.ncbi.nlm.nih.gov/pubmed/19167509.

Yufeng Zang, Tianzi Jiang, Yingli Lu, Yong He, and Lixia Tian. Regional homogeneity approach to fmri data analysis. Neuroimage, 22(1):394–400, 2004.

Y. Zhang, N. Vaidya, U. Iyengar, E. Sharma, B. Holla, C. K. Ahuja, G. J. Barker, D. Basu, R. D. Bharath, A. Chakrabarti, S. Desrivieres, P. Elliott, G. Fernandes, A. Gourisankar, J. Heron, M. Hickman, P. Jacob, S. Jain, D. Jayarajan, K. Kalyanram, K. Kartik, M. Krishna, G. Krishnaveni, K. Kumar, K. Kumaran, R. Kuriyan, P. Murthy, D. P. Orfanos, M. Purushottam, M. Rangaswamy, S. S. Kupard, L. Singh, R. Singh, B. N. Subodh, K. Thennarasu, M. Toledano, M. Varghese, V. Benegal, G. Schumann, and Veda consortium c. The consortium on vulnerability to externalizing disorders and addictions (c-veda): an accelerated longitudinal cohort of children and adolescents in india. Mol Psychiatry, 2020. ISSN 1476-5578 (Electronic) 1359-4184 (Linking). doi: 10.1038/s41380-020-0656-1. URL https://www.ncbi.nlm.nih.gov/pubmed/32203154.

